# Deafness and Sign Language Experience Shift Visual Category Representations

**DOI:** 10.64898/2026.06.09.731097

**Authors:** Edan Daniel Hertz, Jesse Gomez

## Abstract

Childhood visual experiences shape our ability to rapidly recognize faces and objects. Across development, regions processing visual categories that lose relevance, such as hands, are recycled for others. How would visual cortex accommodate a childhood in which hands maintain significance? Through functional magnetic resonance imaging, we demonstrate that high-level visual cortex in Deaf signers develops a unique topography to accommodate the learning of sign language in childhood, with distinct but significant changes observed in hearing signers who acquired sign language in adulthood. These data suggest a new framework for human visual cortex in which the location of regions is not as fixed as once thought, and sociolinguistic experience outside the childhood plasticity period may be sufficient to alter the function of high-level visual cortex.

## Main Text

How does the human brain adapt to support our unique visual requirements? Within a fraction of a second, our visual system can parse a complex visual scene into distinct, comprehensible objects. This outstanding ability is not innate, but rather emerges in response to visual experience during a critical window of childhood plasticity (*1*–*4*), resulting in clusters of neurons in ventral temporal (VTC) and lateral occipitotemporal cortex (LOTC) which support the perception of high-level, ecological stimulus categories such as faces, bodies, words, and buildings. These regions emerge in consistent, anatomically predictable locations (*5*) likely as result of shared viewing patterns interacting with consistent visual features shared by stimuli within a given category (*2, 4, 6*).

Experience is necessary for the emergence of these categories: In illiterate adults lacking visual experience with words (*7, 8*) and non-human primates lacking experience seeing faces (*9*), selective representations for these categories fail to develop in high-level visual cortex. Prior developmental work has shown that unequal ecological relevance across stimulus categories can result in cortical recycling. For example, face- and word-selective representations in the occipitotemporal sulcus (OTS) grow in surface area across childhood and overtake the representation of limbs (*10*) whose relevance as a visual category declines with age. Our certainty that these category-selective regions are spatially consistent and undergo recycling during a critical childhood window are based on findings from humans with typical sensory experiences. This begs the question: how does cortex accommodate an experience in which there is increased demand for processing all visual categories, including limbs? Here, in a group of individuals with unique developmental visual experience, we demonstrate two new principles: the spatial localization of category-selective regions is not purely innate but subject to experience, and the notion that visual experience must occur during childhood for shaping high-level visual representations may have exceptions.

While there is variability in the timing and nature of acquisition, many individuals in the Deaf community learned during childhood to communicate using manual signs. In the United States, American Sign Language (ASL) is a complete, visual language used by both the Deaf and hearing, relying heavily on the use of the upper limbs to convey meaning. Deaf individuals experience qualitatively similar visual stimuli compared to hearing individuals (e.g., faces, hands, scenes) in their daily lives, yet the way in which they are encouraged to pool visual information across space while communicating with ASL—fixating faces and peripherally attending to upper limbs (*11, 12*)—represents an ideal system for modeling how differences in visuospatial sampling of the environment impacts cortical organization. Indeed, while prior behavioral work has shown that Deaf adults may possess superior peripheral vision (*13*), the neural correlates of these and higher-level visual behaviors are poorly understood from a lack of research on the Deaf visual system.

Here we ask two questions about human brain plasticity: How does the visual system accommodate its development to support a vision-based language? Second, is learning sign language in adulthood, after the plasticity window of childhood, sufficient to induce changes in the functional topography of high-level visual cortex? We combined structural and functional magnetic resonance imaging (MRI) with behavioral metrics to assess how category-selective representations in visual cortex respond to the acquisition of American Sign Language (ASL). We include members of the Deaf community fluent in ASL from early childhood (Deaf Signers, N=11), hearing signers who learned ASL in adulthood for the purpose of becoming state-registered interpreters or teachers (Hearing Signers, N=11, with more than 10 years of ASL experience), and controls with no ASL experience (Controls, N=11). Groups were matched for age. Hearing signers, who acquired sign language expertise later in life, allow us to identify whether the effects of learning sign language on the visual system are specific to childhood or deafness.

We employed a bidirectional video system to enable Deaf participants and interpreters to communicate within the MRI machine (**Fig 1A**). In addition to structural neuroimaging for reconstruction of each participant’s cortical surface, participants completed a visual category localizer (**Fig 1B**) to identify in each individual’s brain the representations for faces, hands, and places in high-level visual cortex, which spans both the ventral temporal and lateral occipitotemporal surfaces (**Fig 1C**). We focus analyses on these three categories given their importance for Deaf signers, who fixate on faces during sign language communication, attend to upper limbs including hands for sign perception, and monitor the visual scene in the absence of audition.

**Figure 1:**
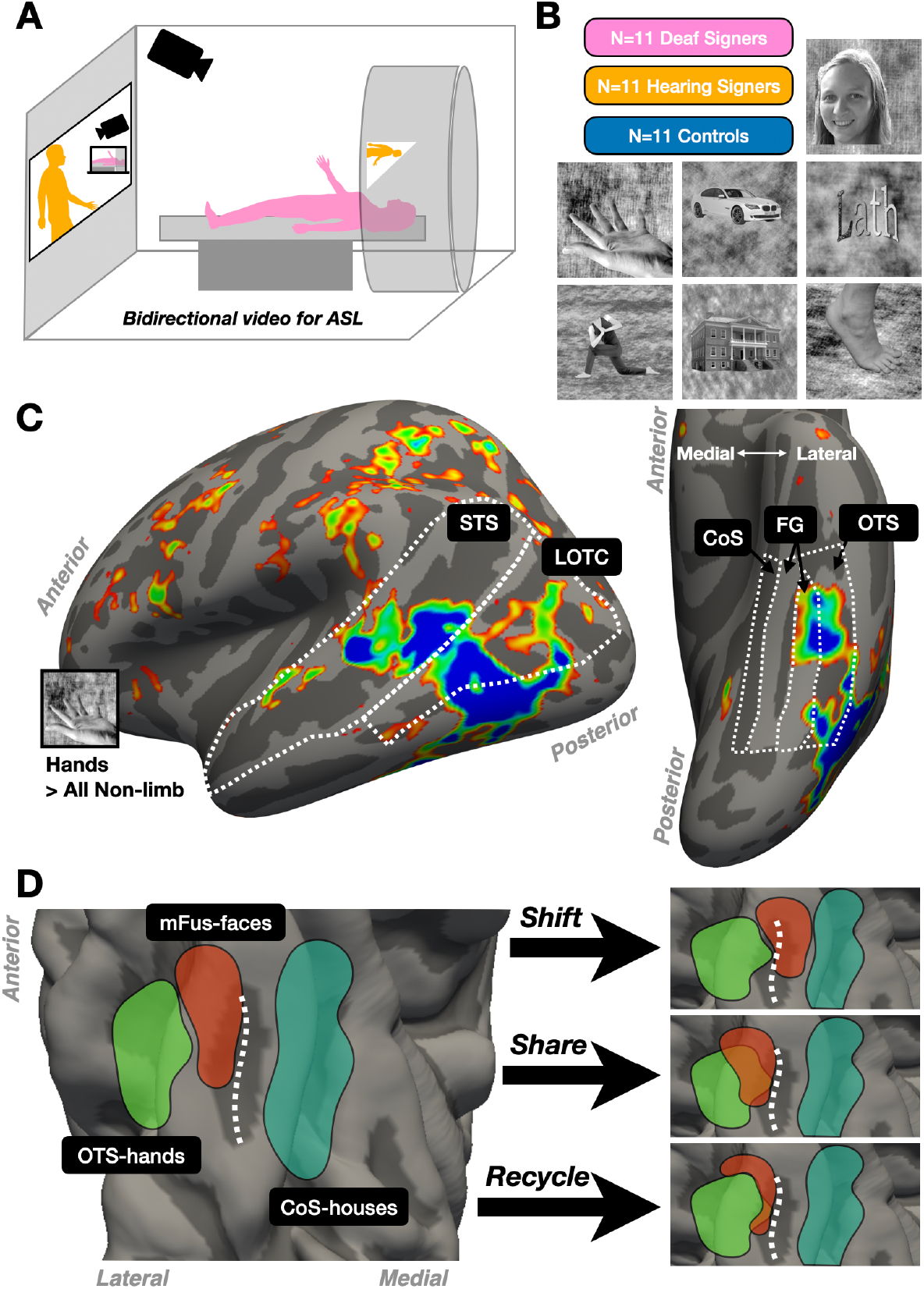
Neuroimaging of high-level visual cortex and theories for how its functional topology accommodates sign language. **(A)** Bidirectional video system to enable Deaf signers to communicate during magnetic resonance imaging. **(B)** Example stimulus categories from the visual category localizer for identifying regions of cortex that preferentially respond to faces, hands, or houses, with remaining stimuli acting as control categories to evaluate selectivity of neural responses. Face image is that of an author for purposes of example. **(C)** Example map depicting hand-selective responses measured with fMRI in a Deaf participant. Vertices are colored by t-value denoting significance of the contrast hand > all other non-limb stimuli, thresholded at a t-value of 3. Anatomical regions of interest (ROI) across lateral and ventral visual cortex are outlined in white (OTS = occipitotemporal sulcus, FG = fusiform gyrus, CoS = collateral sulcus). The lateral visual stream is parcellated into the superior temporal sulcus (STS) and lateral occipitotemporal cortex (LOTC). The ventral visual stream is parcellated in the occipitotemporal sulcus (OTS), the lateral Fusiform gyrus (lat-FG), the medial Fusiform gyrus (m-FG), and the collateral sulcus (CoS). **(D)** Illustration of possible changes in functional topography to accommodate a hypothesized increase in hand representations in response to the learning of sign language.

If the representation for hands is not developmentally pruned in Deaf Signers, how does visual cortex make room for a region that is typically recycled in the hearing? With this data, we will adjudicate between three possible scenarios. First, as previous developmental work has shown, increasing demand for processing the upper limbs in ASL may increase the representations of limbs and “recycle” neighboring representations for faces (**Fig 1D**). A second possibility is that there may be an increase in “shared” cortical representation for faces and limbs. Support for a “share” model of visual topographic change (**Fig 1D**) comes from prior work that has shown that the overlap that can occur between neighboring representations of faces and bodies allows for clutter-tolerant representations of the face, body, and whole person (*14*). A third possibility, which has not yet been empirically observed in human visual cortex, is that category-selective regions may shift their spatial location to make room for increasing representation of limbs in lateral visual cortex. In this “shift” model, for example, neighboring representations such as face-selective cortex on the fusiform gyrus may become more medially located (**Fig 1D**). To test this, we quantify the cortical surface area representing hand, face, and place stimuli in high-level visual cortex spanning LOTC and VTC. Group differences in the size of category representations and their overlap can be used to assess evidence for the “share” and “recycle” models of visual cortex development. To quantify how regions may alter their spatial location in the “shift” model, we focus specifically on VTC given that the spatial locations of face, hand, and place-selective regions have been thoroughly characterized, providing a strong *a priori* model of typical high-level vision.

### Sign language expertise increases visual representation for hands

We first focus on hand stimuli, for which there are clustered representations in both the lateral and ventral visual streams (*15*). We performed a bootstrapping analysis (1000 iterations, two-tailed CI = 95%) to quantify the cortical surface area responsive to hands in both Deaf Signers and Hearing Signers compared to Controls. Deaf Signers demonstrated a significant increase in hand-selective cortical surface area (+372 vertices, CI = [158 573], p < 0.0001, Cohen’s d=1.71) in the left hemisphere of ventral temporal cortex (**Fig 2a**). The right hemisphere showed a numerically larger surface area as well (+152 vertices, p = 0.27, cohen’s d=0.51, **Fig S1a**). Hearing Signers showed a smaller but significant increase of 354 vertices in the left hemisphere of ventral temporal cortex (CI = [7 720], p = 0.044, Cohen’s d=0.90) compared to controls (**Fig 2a**), and a similar effect on the right (+294, CI [-13 610], p = 0.066, Cohen’s d = 0.86). On the lateral surface, we find a robust increase in hand selectivity only amongst Deaf Signers in the right hemisphere, with a median increase of 1284 vertices (CI = [383 2270], p = 0.006, Cohen’s d = 1.16) but not the left hemisphere (p = 0.19, Cohen’s d = 0.58), and no significant change in surface area within Hearing Signers in either hemisphere (left: p = 0.32, Cohen’s d = 0.44; right: p = 0.25, Cohen’s d = 0.47; **Figs 2b, S1b**).

**Figure 2:**
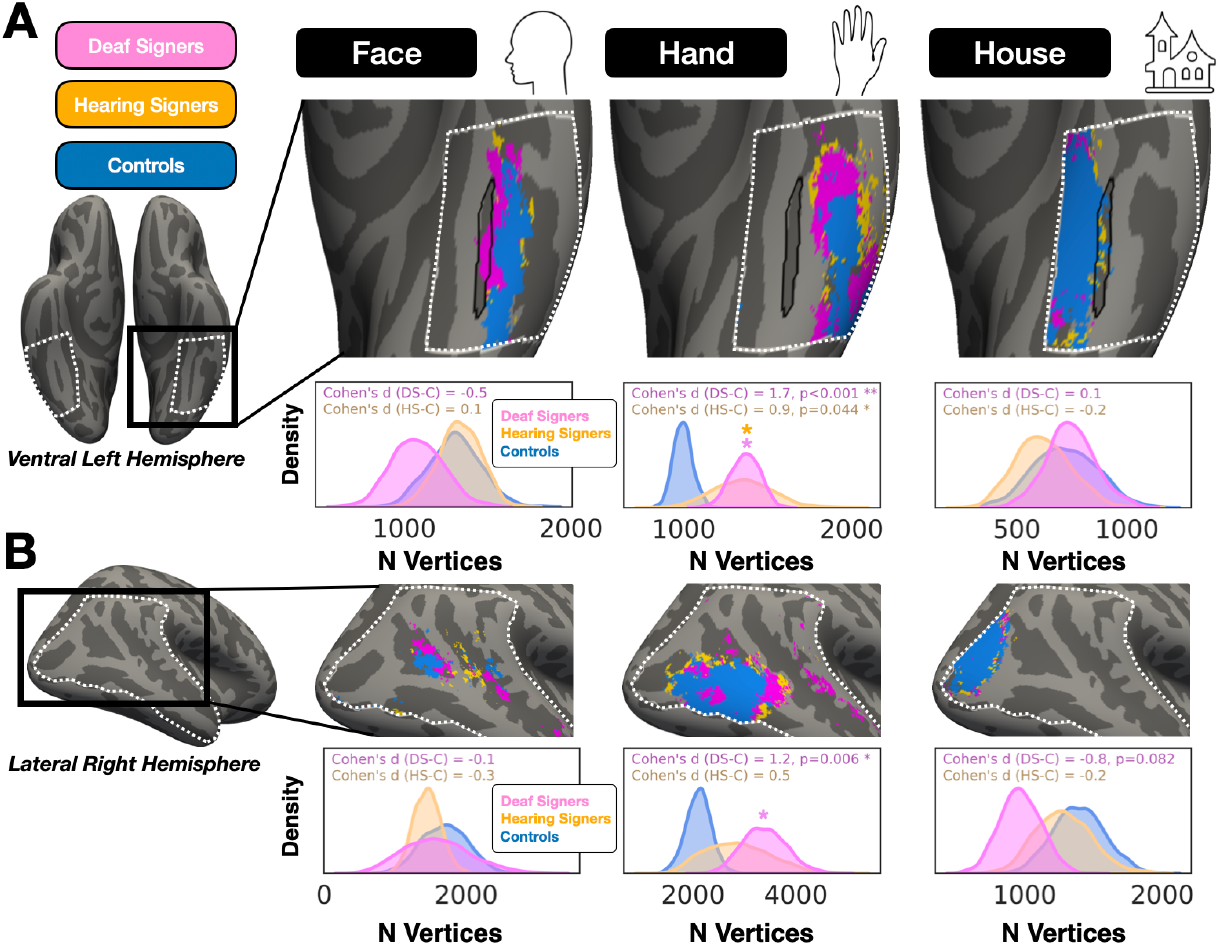
Expansion of hand representations in Deaf and hearing signers. **(A)** Face, hand, and house selective regions in ventral temporal cortex (dashed white border). Contrast maps were thresholded at T>3 and binarized within individuals, then summed across individuals per group and thresholded at minimum of 4 participants to create the group-level category selective region. The mid-Fusiform sulcus (MFS) is outlined in black as an anatomical reference. Histograms depict boot strapped (n = 1000) distributions of surface area (number of vertices in average cortical space) in each group, drawn with replacement. **(B)** Category-selective regions of similar categories but within lateral visual cortex (dashed white border).

Is areal expansion for the visual representation of hands also observed for the categories of faces and places? The surface area of face-selective cortex on the ventral and lateral cortical surfaces was not significantly different in either Deaf or Hearing Signers compared to Controls (p’s ≥ 0.4, **Fig 2a-b, S1a-b**). The same was true for the representation of places in ventral temporal cortex. In lateral occipitotemporal cortex, Deaf Signers demonstrated smaller cortical surface area responsive to place stimuli but this difference was only significant in the left hemisphere (L eft: median difference = -559 vertices, CI = [-930 -174], p = 0.004, Cohen’s d = -1.29; Right: -420, CI = [-891 46], p = 0.08, Cohen’s d = -0.84; **Fig S1**).

### Does increasing representation for hands result in shared cortical representations?

Face and limb representations neighbor in both ventral and lateral visual cortex (*16*). This pattern of proximal representations has been suggested as an organization principle in the visual cortex, and has been shown to be an efficient way to computationally implement clutter-tolerant representations of the face, body, and whole person (*14*). With the increase in information coming from hands during signed conversations, there are two possible outcomes. First, the increased relevance of hands may drive more specificity in their representation, with a more distinct border separating hand from non-hand stimuli. Second, the need to simultaneously process hand and face-relevant information during sign language may result in more cortical sheet which shares responsiveness to both stimuli (**Fig 1d**). Given that face and limb regions border one another and shared cortical representation could in theory occur in either direction–medially into the occipitotemporal sulcus (OTS) or laterally into the fusiform gyrus (FG)–we anatomically parcellate visual cortex to spatially quantify any effect into the medial FG, lateral FG, and OTS.

In each group we compute the conjunction between face and hand selectivity within VTC. While we find overlapping representations within each group of statistically equivalent size (**Fig 3a**), the location of this dual representation for faces and hands is distinct across groups when examining where across the FG and the OTS this overlap lies. Control and Hearing Signers show a majority of face-hand overlap within the OTS while Deaf Signers show more overlap in the lateral FG in both hemispheres (**Fig 3a**). In a Linear Mixed Effects Model (Normalized N Vertices ∼ Group + ROI +

**Figure 3:**
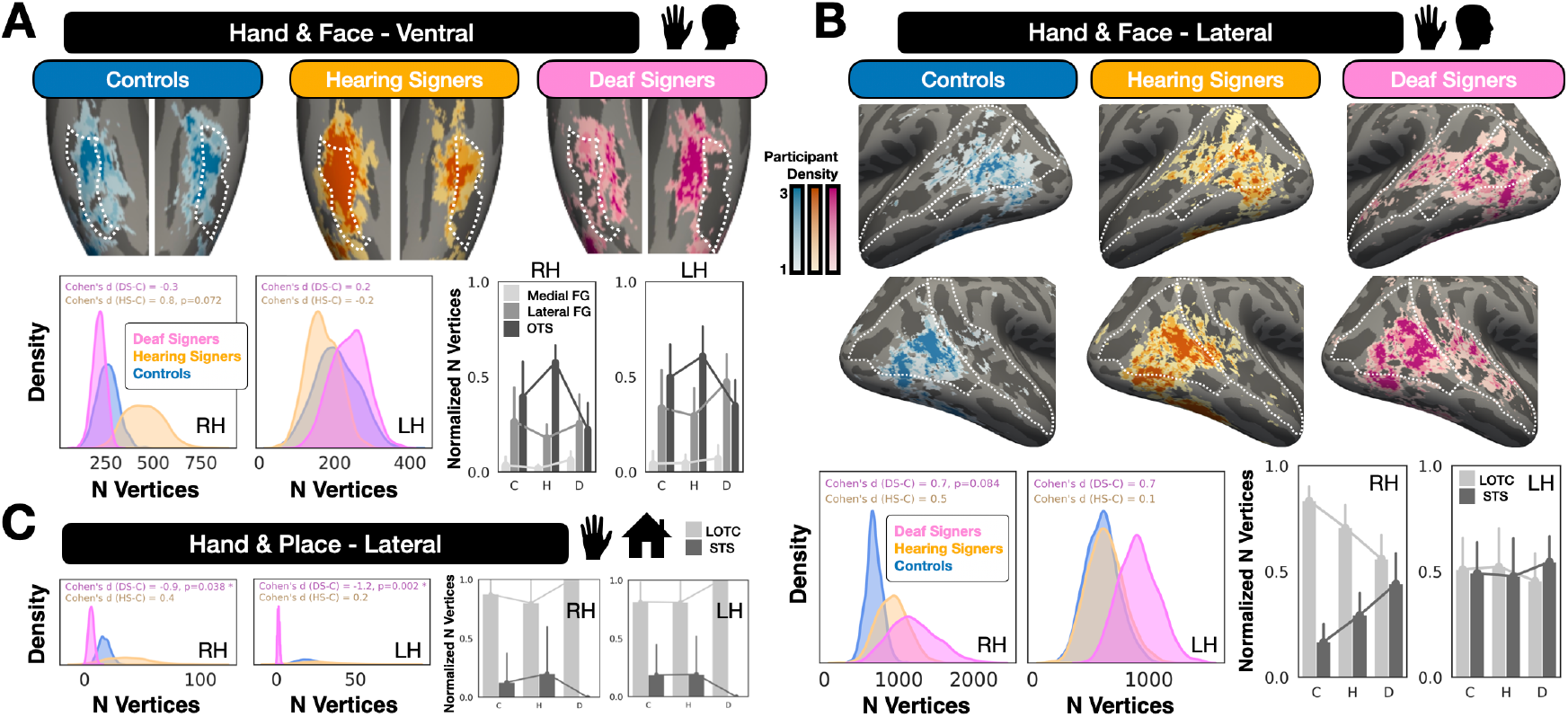
Location of overlap in category representations is distinct across groups in ventral and lateral visual cortex. **(A)** Upper row: The overlap of hand and face selectivity in bilateral ventral temporal cortex shown on the inflated average cortical surface. The OTS is outlined in dotted white for reference. Selectivity for hands and faces was computed as the preferred versus inanimate stimulus categories, so that the contrast did not bias against overlap. Overlap area is the number of vertices within the conjunction of hand and face selectivity maps within the anatomical ROI. Vertices of overlap are binarized in each participant then summed in cortical average space, with colormaps denoting participant density. Right hemisphere is shown on left. Lower row: Histograms depict bootstrapped distributions of overlap area. Skyline bar plots depict the proportion of vertices within each participant group for each anatomical subregion (normalized within-participant by number of vertices in a participant’s hemisphere). RH/LH= right/left hemisphere. **(B)** Similar to panel A but for lateral visual cortex, with the superior temporal sulcus (STS) and lateral occipital temporal cortex (LOTC) outlined in dotted white for reference. **(C)** Similar to previous panels for the overlap of hand and place selectivity in lateral visual cortex where hand and place regions share a border.

Group*ROI + (1|Sub)), we find a significant Group x ROI interaction in the right hemisphere (χ_2_ = 16.8, p = 0.002), with a marginal effect on the left (p = 0.066). The significant interaction on the right was specifically driven by the face-hand overlap on the OTS, where Deaf Signers show significantly less overlap compared to controls (β = -0.33, p = 0.027, CI [-0.6 -0.04]), and Hearing signers show an opposite trend, with an increase in OTS face-hand overlap compared to Controls (β = +0.26, p = 0.08, CI [-0.03 0.6]).

On the lateral surface, we find similar evidence which further distinguishes Deaf Signers from Hearing Signers and Controls. Dividing the lateral surface into occipitotemporal cortex (LOTC) and the superior temporal sulcus (STS), we find that groups significantly differ in the overlap location of their face-hand representations (**Fig 3b**). In the right hemisphere, the overlap between face and hand representations is evenly distributed across LOTC and STS in deaf signers, while in Hearing Signers and Controls the majority of this overlap occurs in LOTC (Linear Mixed Effects Model Group-ROI interaction: χ_2_ = 23, p < 0.0001). While both Hearing Signers and Deaf Signers showed less LOTC dominance and more face-hand overlap on the STS, the effect observed among Deaf Signers (β = +0.55, p <0.0001, CI [0.3 0.8]) was twice as large as the moderate spatial modulation observed in Hearing Signers (β = +0.25, p = 0.03, CI [0.02 0.5]), when both groups were compared to Controls. In the left hemisphere, while Deaf Signers show numerically more vertices dually tuned for faces and hands in the STS, the distributions across STS and LOTC is statistically equivalent across all groups (Group-ROI interaction: p = 0.7).

Unlike ventral temporal cortex, LOTC contains a place selective region in the lateral occipital sulcus (LOS-places) that abuts the limb representations of the lateral visual stream (*17*). To investigate the possibility that hand representations interact with place selectivity in LOTC, we performed a similar analysis quantifying the spatial arrangement of place-hand overlap. Strikingly, both Hearing Signers and Controls show overlap between place and hand representations while Deaf Signers show virtually none (**Fig 3c**; Deaf Signers to Controls bootstrap analysis; Left: median difference = -18 vertices, CI = [-31 -6], p = 0.002, Cohen’s d = -1.25; Right: median difference = -11 vertices, CI = [-22 -1], p = 0.038, Cohen’s d = -0.94). In a Linear Mixed Effects Model comparing the subregions, we find no Group x ROI interaction on either hemisphere (Left: p=0.6 ; Right: p=0.3), and no main effect for group (Left: p=0.8 ; Right: p=0.6).

### Sign language experience alters the spatial topography of high-level visual cortex

While there is shared representation for faces and hands in all groups, it appears that the location of where this sharing occurs on the cortical sheet is uniquely medial in Deaf signers (**Fig 3a**). Has the spatial location of category representations shifted to accommodate the learning of sign language? To quantify this, we take a three-pronged approach and focus on VTC where the position of category-selective regions is well understood. First, we ask how regions are positioned relative to anatomical landmarks. In VTC, we quantify the distribution of face-selective surface area across anatomically-defined regions: medial FG, lateral FG, and the OTS. Groups significantly differ in how face-selectivity is distributed across VTC, with Deaf Signers showing a higher percentage of face-selectivity residing on the medial FG compared to the OTS (**Fig 4a, S2a**). For house selectivity located more medially, we focus on lateral FG, medial FG, and the collateral sulcus (CoS). We also find a significant group by ROI interaction, with house selectivity in Deaf Signers also shifting medially, with more selectivity within the Collateral Sulcus (CoS) and less on the medial FG compared to other groups (**Fig 4a, S2a**). For hand selectivity, lateral visual cortex shows the most striking group differences. Quantifying the distribution of hand selectivity across LOTC and the STS in the right hemisphere, Deaf Signers show an even distribution of hand representations across these two structures, while Controls and Hearing Signers show relatively more hand selectivity in LOTC (**Fig 4a, S2b**).

**Figure 4:**
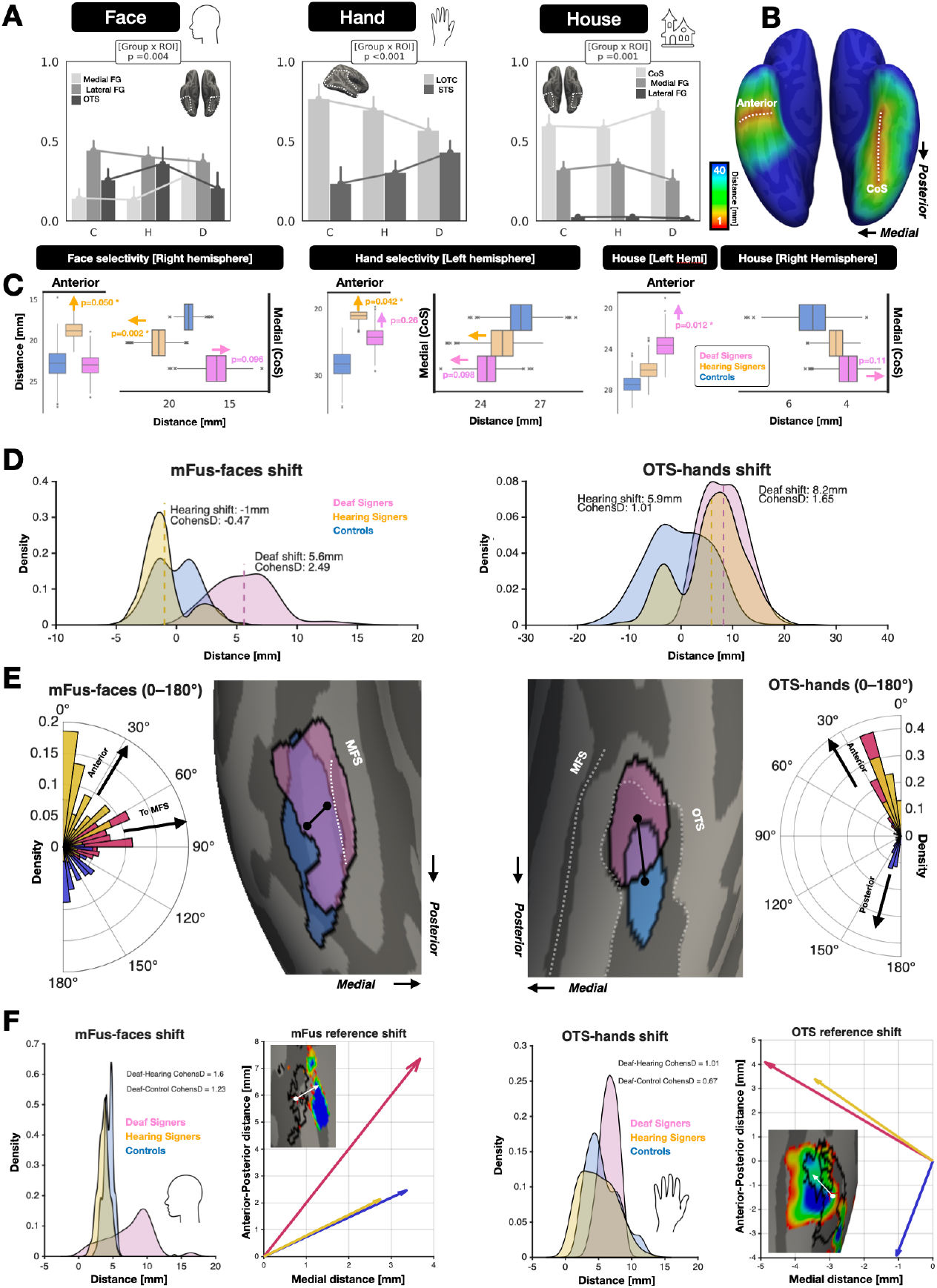
Location of hand and face representations shift in Deaf signers. **(A)** Normalized bilateral surface area of face, hand, and house selectivity in each participant group as a function of anatomical region of interest. C = Controls, H = Hearing Signers, D = Deaf Signers. Error bars denote bootstrapped 95% confidence intervals around the group means (n = 1000). **(B)** Distance map on the FreeSurfer average brain denoting the minimum distance in millimeters of a given vertex to vertices within the fundus of the collateral sulcus (CoS) used for deriving the mean shift of functionally-defined ROIs in ventral temporal cortex. Line drawn at anterior FG used for anterior distance measurements is also illustrated in dotted white. **(C)** Box plots depicting bootstrapped mean distance values of face-selective vertices on the right FG, hand-selective vertices in the left OTS, and house-selective vertices in the CoS from either the anterior or medial anatomical reference point shown in panel B. **(D)** Histogram of bootstrapped distances of the centroid of the right mFus-faces or the left OTS-hands ROI. Distance measured relative to Control group centroid. **(E)** Polar histograms representing the angle of centroid displacement of group-average right mFus-faces or left OTS-hands ROIs relative to a vector that lies parallel with the mid-Fusiform sulcus (MFS). Inflated cortical surface zoomed on ventral temporal cortex shows an example bootstrapped group-average mFus-faces (left) and OTS-hands (right) for Control participants (blue) and Deaf Signers (pink). **(F)** Distance histograms similar to panel D but depicting bootstrapped centroid distances of each group-average ROI relative to an atlas-based definition of the face and hand ROI. Inset cortical surfaces show an example mFus-faces (left) and OTS-hands (right) from a Deaf Signer with the atlas-based definitions of mFus-faces and OTS-bodies outlined in black for reference.

We can further quantify movement with a complementary analysis, asking for a given category-selective region how far in millimeters its vertices are on average from an anatomical reference. We thus derived distance maps from two perpendicular anatomical structures to quantify medial-lateral, and anterior-posterior movement respectively from the CoS and the anterior FG (**Fig 4b**). Similar to the anatomical ROI results, face-selective cortex appears to shift in the medial-lateral direction, with Deaf and Hearing signers showing distinct shifts relative to Controls. Deaf Signers demonstrate a more medially-located face-selective region (**Fig 4c**). Face-selective vertices are also on average more anterior and lateral in Hearing Signers compared to other groups. For hand-selective cortex, Deaf and Hearing Signers show selectivity that is located more medially and anteriorly compared to Controls. Further replicating the anatomical ROI analysis, house selectivity is relatively more medially located in Deaf Signers. It is also more anterior in Deaf Signers compared to Controls (**Fig 4c**). Other hemispheres show similar effects (**Fig S3**).

### Shifting centroids of face and hand regions in ventral temporal cortex of signers

To complement this analysis with a second approach that is more sensitive to the centroid movement of functional representations, we perform a surface-based analysis in which average category-selective regions are bootstrapped (n=500) from within each group and vectors are derived describing the distance between centroids (weighted center vertex) of Deaf or Hearing Signers relative to Controls. On each iteration, a second Control ROI is produced to evaluate variability of the Control group to themselves. We focus on the face-selective region of the middle fusiform gyrus (mFus-faces) and the hand-selective region located just laterally in the OTS (OTS-hands). We quantify both the length and angle of functional displacement vectors. For OTS-hands, Deaf Signers show a mean shift of 8mm relative to the centroid of controls (**Fig 4d**) at an angle of roughly 27-degrees (**Fig 4e**) from an axis parallel with the MFS (CI = [2 15], Cohen’s D = 1.65). Hearing Signers shift 5.9mm in a similar direction (CI = [-5.1 14.9], Cohen’s D = 1.01, **Fig 4d-e**), while Controls shift around themselves equally in the anterior and posterior direction. For face-selective cortex, the average mFus-faces of Deaf Signers has shifted medially 5.6mm relative to the centroid of controls (CI = [1.06 12.56], Cohen’s D = 2.49, **Fig 4d**) at an angle of 85-degrees relative to the MFS (**Fig 4e**), nearly perpendicular to the average displacement vectors in Hearing Signers and Controls. The mFus-faces centroid of Hearing Signers does not show this medial movement, shifting on average 1mm relative to controls in a direction distinct from Deaf Signers (CI = [-3.7 3.75], Cohen’s D = -0.47), largely overlapping with that of controls and showing displacement vectors oriented anteriorly.

### Hand representations specifically expand outside of an independent probabilistic region

Lastly, we can compare face- and hand-selective ROIs to probabilistic definitions of these regions derived from independent data. We compared our selectivity maps to the OTS-body and the Fus-faces regions defined in the visfAtlas, a functional atlas defining the location of category-selective regions in human occipito-temporal cortex (*18*). While the OTS-body region was defined using stimuli that contained other limbs and bodies, it remains a useful independent reference to which we can compare all three groups. For each participant, we quantified the number of vertices included within, and outside of, the expected region of the OTS-body and the Fus-faces region. For hand selectivity, we find that both Deaf and Hearing signers show a significantly stronger deviation from the reference OTS-body region compared to controls, driven by an increase in the vertices outside of the independent region (Linear Mixed Effect Model: Vertices Outside ∼ Group + Hemisphere + Group*Hemisphere + (1 | Sub_ID), Deaf versus Controls: β = +315, p = 0.015, CI [60 570]; Hearing versus Controls: β = +280, p = 0.03, CI [26 535]; Main Effect of Group: χ_2_ = 7.1, p=0.029). In contrast, the area of intersection with the reference region remained unchanged (Deaf versus Controls: β=+27, p=0.6; Hearing versus Controls: β=+62, p=0.19). This indicates an additive increase of outside-of-reference selectivity. That is, the observed increase in hand representations in the experimental groups is not a general effect with random topography, but a spatially specific expansion that occurs outside the borders of the reference ROI. For the Fus-faces region, we find no group differences in the number of vertices overlapping the expected functional region or outside of it (p’s > 0.45).

We can also repeat our displacement vector analysis using the centroid of visfAtlas regions as reference points to ask if Deaf and Hearing Signers show unique spatial deviation from an independent group of participants. For both hand and face-selectivity, all groups show significant displacement from the reference ROI (confidence intervals exclude 0), but show distinct effect sizes and movement directions. For OTS-hands, the average hand selectivity of Deaf and Hearing Signers is displaced medially relative to the centroid of the reference ROI while the displacement vector in Controls points posteriorly (**Fig 4f**). Deaf Signers show the largest shift relative to both Hearing Signers (Cohen’s D = 1.01) and Controls (Cohen’s D = 0.67). We also replicate the displacement effect for face-selective cortex: the average mFus-faces ROI in Deaf Signers is shifted medially relative to the reference ROI compared to both Hearing Signers (Cohen’s D = 1.6) and Controls (Cohen’s D = 1.23), oriented more medially towards the anterior tip of the MFS compared to both groups (**Fig 4f**).

### Changes in the symmetry of visual responses across hemispheres of the brain mirror viewing patterns

Learning sign language appears sufficient to alter the local spatial arrangement of high-level representations in visual cortex. Is the broader distribution of visual processing across lobes and hemispheres of the brain also responsive to sign language experience? A hallmark of adult high-level visual cortex is that representations for faces are not evenly distributed across the cerebral hemispheres, with larger representations in the right hemisphere (*19*) thought to result in part from the left lateralization of language processing (*20*). Because Deaf Signers perceive language in a unique, unimodal fashion in which visual information from faces and hands is heavily used for social communication, we were interested in exploring how sign language experience might affect the hemispheric biases in representation of the different visual categories. To this end, we calculate a laterality bias in the cortical surface area selective for a given stimulus category as the ratio of left hemisphere surface area normalized by the sum of left and right (**Fig 5a**). In particular, prior work sets a strong a priori hypothesis that groups should show a rightward lateralization bias for the processing of faces.

**Figure 5:**
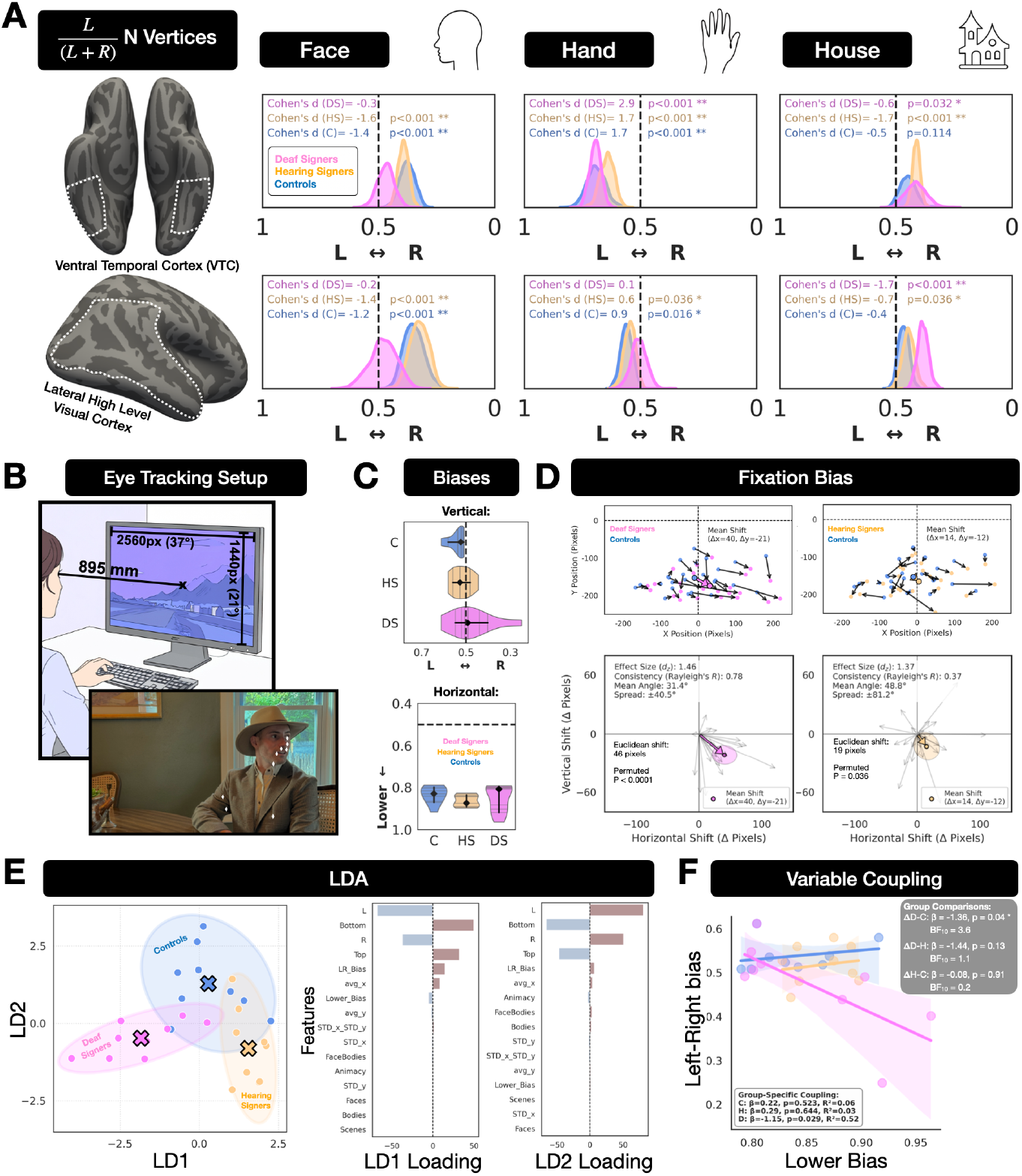
Lateralization shifts in Category Representations and Viewing Biases. **(A)** Lateralization of category representations on the Ventral (Top row) and Lateral (Bottom row) surfaces. Left to Right Ratio is calculated within-participants for each visual category (face, hand, house) and visual stream (ventral/lateral), and defined as the number of vertices on the Left hemisphere divided by the sum of vertices on both Left and Right, such that Ratio >0.5 indicates a leftward bias. Histograms depict bootstrapped distributions of L/R Bias. **(B)** Experimental Setup for Eye Tracking Experiment. Top image generated by Adobe Firefly. Bottom inset image shows example movie frame from the movie stimuli shown to participants, where white markers denote example fixation points of participants. Actual image is a reenactment photo depicting an author for illustration purposes. **(C)** Viewing Biases across the full experiment, in the vertical and horizontal dimensions. Ratio = 0.5 indicates no Bias. **(D)** Group differences in fixation position within each movie clip. Left: Deaf Signers vs Controls. Right: Hearing Signers vs Controls. Top row: Each vector connects the fixation centroids of the Control (start) and Signer group (end) for a single stimulus (N=25 stimuli), in the original screen space (pixels). Dashed lines mark the screen center. Bottom Row: similar to top row, but centralizing Control group centroids at the origin (0,0) to test the consistency of direction of displacement. Colored arrow depicts the average vector across all stimuli, indicating the direction of fixation bias. Colored shaded area depicts the 2D 95% confidence ellipse around the group average displacement. **(E)** Linear Discriminant Analysis (LDA) testing group separability based on eyetracking features alone. Left: Individual data points (participants, N=9 per group) plotted against the two LD components. Bar plots show feature loadings ranked by absolute value. **(F)** Deaf signers show a unique relationship between L-R Bias and Lower Visual Field Bias.

### Deaf Signers show no rightward bias in face-selective representations

Across ventral and lateral visual cortex, we find that the well-characterized rightward bias for face representations is surprisingly absent or reversed in Deaf Signers (**Fig 5a**). In both the ventral and lateral surfaces, there is a significant difference in the lateralization of surface area selective for faces (Linear Mixed Effects Model: Ratio ∼ Surface + Group + Surface*Group +(1|Sub), Main effect for Group: χ_2_ = 8.32, p = 0.016). This effect was specifically driven by a stronger rightward bias in Deaf Signers compared to controls (β = 0.135, p = 0.02), with no difference between Hearing signers and Controls (p = 0.7), and no Surface-Group interaction (p = 0.45). In a Bootstrapping analysis, both Hearing Signers and Controls show significant rightward lateralization for faces in ventral (p < 0.0001) and lateral (p ≤ 0.002) surfaces while Deaf Signers do not (p ≥ 0.4). For hand representations, Hearing Signers and Controls both show a significant leftward bias in lateral visual cortex (HS: median ratio = 0.54, p = 0.036, CI [0.503 0.597], Cohen’s d = 0.63, and Controls: median ratio = 0.56, p = 0.016, CI [0.51 0.61], Cohen’s d = 0.87) while Deaf Signers do not (p = 0.8, CI [0.44 0.57], Cohen’s d = 0.08). In the ventral surface, all groups show a strong leftward bias for hand representations (p < 0.0001, median ratios ≥ 0.64, Cohen’s d’s ≥ 1.67). For place representations, we find that both Hearing and Deaf Signers have significant rightward biases, while Controls do not. Deaf Signers have a strong and significant rightward bias in place representations on the lateral surface (median Ratio = 0.39, p < 0.0001, CI [0.34 0.43], Cohen’s d = -1.7), significantly stronger than Controls (median difference = -0.08, CI [-0.15 -0.02], p = 0.004), with Hearing Signers showing a similar, albeit weaker rightward bias (median Ratio = 0.45, p = 0.036, CI [0.39 0.496], Cohen’s d = -0.71; **Fig 5a**). Interestingly, on the ventral surface, both Hearing and Deaf Signers exhibited a significant rightward bias in place representations (HS: median ratio = 0.41, p < 0.0001, CI [0.38 0.45], Cohen’s d = -1.7; DS: median ratio =0.41, p = 0.032, CI [0.33 0.49], Cohen’s d = -0.63), potentially driven by the expansion of hand representation on the ventral left hemisphere (**Fig 2a**). Overall, Deaf Signers demonstrate a unique hemispheric symmetry in their representation of faces and hands in the ventral and especially lateral visual streams; a surprising observation given the strong precedent set by prior neuroimaging work documenting robust rightward biases in the visual processing of faces.

### Unique Viewing Patterns for Dynamic Social Stimuli

The lateral visual stream in humans shows preferential responses to dynamic (*21, 22*) and social stimuli, leading to the hypothesis that it is a distinct network specialized for social perception (*23, 24*). The symmetric representation of face and hand categories in Deaf Signers in lateral visual cortex suggests that the way in which they visually sample dynamic social stimuli may be unique. Maturing face perception abilities are typically associated with a decrease in an initial leftward childhood bias for viewing faces which becomes increasingly central into adulthood. This behavioral change is coupled with increasing representation of central visual space in face-selective regions as measured with receptive field mapping (*6*). This prior work provides a direct link between the representation of ecological visual stimuli and real-world viewing behaviors, and suggests that sign language acquisition may sculpt viewing behaviors of social scenes involving faces and limbs. To test the hypothesis that Deaf Signers show distinct visual sampling of social stimuli, the same participants who completed neuroimaging were eyetracked while free-viewing movie stimuli (**Fig 5b**). They watched 25, 30-second clips from Hollywood movies, specifically selected for socially enriched scenes (*25*). Critically, to maximize the perceptual experience across groups, no sound was included, and eyetracking occurred outside the scanner. Movie stimuli were downsampled to 150 frames (2Hz), and labeled such that pixels containing faces, bodies, or background scenes could be differentiated (*26*).

For each frame, we quantified: the percentage of total fixations for each object category, relative percentage of face versus body viewing, relative percentage of animate versus inanimate viewing, the raw number of fixations made in each half of the stimulus frame (left, right, upper, lower), the average x and y positions, standard deviations on the horizontal (x) and vertical (y) axes, and a measure of spatial viewing bias defined as the ratio of the number of fixations made to the left side of the visual stimulus normalized by the number of fixations to the left and right, as well as a similar bias metric for upper and lower visual space. To test the hypothesis that symmetric representations across hemispheres of the lateral visual stream in Deaf Signers results in symmetric viewing patterns, we first compare the left-right ratio of viewing in each group. Interestingly every Control participant demonstrated a leftward viewing bias (bootstrapped p < 0.0001, median = 0.54, CI [0.52 0.56], Cohen’s d = 3.25), while Deaf Signers show statistically symmetric viewing patterns with a slight bias towards the right visual field (median = 0.48, p = 0.44, Cohen’s d = -0.83, **Fig 5c**). Hearing Signers show an intermediate effect with a numerically leftward viewing bias but overlapping the zero-bias value as a group (median = 0.54, p = 0.19, Cohen’s d = 1.3). All groups also demonstrated a significant lower visual field viewing bias (medians > 0.81, p < 0.0001, **Fig 5c**). To quantify the consistency of these effects across individual stimuli (30-s movie clips), we computed the centroid of fixation (mean x,y coordinates) averaged across all frames and participants within a group and derived vectors describing the directional difference between Controls and each signing group. When visualizing these vectors, we find a striking group difference in which Deaf participants show a significantly greater fixation bias towards both a rightward and lower point of visual space (**Fig 5d**) relative to Controls (Cohen’s D = 1.46, Permutation p < 0.0001, Δx = +40, Δy = -21) for nearly every stimulus. When centralizing vectors relative to the mean Control centroid, Deaf Signers show a clear preference for sampling visual space distinctly towards the right and lower visual field (**Fig 5d**). Hearing Signers also show significantly distinct viewing patterns relative to Controls, albeit less striking (Cohen’s D = 1.37, Permutation p = 0.036, Δx = +14, Δy = -12) (**Fig 5d**).

To evaluate all viewing features holistically, we employ a data-driven machine learning approach to ask if groups can be distinguished based on their viewing patterns. Fixation metrics were entered into a linear discriminant analysis (LDA) which seeks to identify a linear combination of features to separate groups. When scattering participants against the two linear discriminant (LD) dimensions, Deaf Signers, Hearing Signers, and Controls group into distinct clusters with the mean coordinate of Control and Deaf Signer groups lying ±2 standard deviations outside the other groups’ distribution, while the mean coordinate for Hearing Signers was located within the distribution of Controls (**Fig 5e**). Cluster validity, quantified by a silhouette coefficient of 0.29, indicated moderate class separation with mean inter-cluster exceeding intra-cluster distance. The top features contributing to group separation were the number of fixations to the left, right, top, and bottom visual field, as well as their bias metrics and average x position (**Fig 5e**). Lastly, given the importance of the left-right and lower viewing biases for distinguishing group viewing behavior, we asked if the coupling between these two features is significantly different between groups. When evaluating the linear relationship between horizontal and vertical viewing biases (**Fig 5f**), we find that only Deaf Signers demonstrate a significant and negative correlation between the two viewing metrics (β = -1.1, R_2_ = 0.52, p = 0.03). This relationship is also significantly different from the linear relationship in the other two groups (Deaf v. Control: **Δ**β = -1.36, p = 0.04, Bayes Factor = 3.6; Deaf v. Hearing: **Δ**β = -1.44, p = 0.13, Bayes factor = 1.1). Hearing Signers and Controls showed no significant differences between one another (**Δ**β = -0.08, p = 0.91, Bayes Factor = 0.2).

## Conclusions

Together, these results suggest that the acquisition of sign language expertise in both Deaf individuals and hearing interpreters who did not acquire sign language until early adulthood, is sufficient to alter the spatial topography and viewing patterns of ecologically relevant stimuli such as faces and hands. This finding suggests that the idea of a critical window in vision should be readdressed, as the current results are at odds with the idea that specialized category-selective regions are responsive to visual experiences exclusively during childhood (*4, 9, 10, 27*). The uniquely dramatic expansion of hand-selective representations in the lateral visual cortex of Deaf Signers is consistent with recent work proposing a lateral visual stream specialized for social perception in humans (*24*). The surprising lack of a rightward hemispheric bias for face representation in Deaf Signers is consistent with theories positing that the left lateralization of language drives the right lateralization of face processing (*20*); when faces are integral for language, as in ASL, their increased sociolinguistic relevance may lead to bilateral representation. The recent description of somatosensory representations of the body across lateral visual cortex (*28*) implies many of the current findings could be driven by additional changes in the visuomotor representation of effectors for the face and hand in parietal and prefrontal cortex. Overall, these results demonstrate an expanded framework of plasticity in human high-level visual cortex which is capable of accommodating an additional visual language. These data strongly support previous findings indicating that deaf and hard-of-hearing children with early access to sign language benefit cognitively (*29*) by showing that high-level visual cortex can readily incorporate expanded representations for hand stimuli integral to signing.

## Acknowledgments

We are grateful for the support of the sign language experts and members of the Deaf community who participated, and for the research infrastructure and technical assistance from the Scully Brain Imaging Center of the Princeton Neuroscience Institute. We are also grateful to Patricia Hoyos, Federico d’Oleire Uquillas, Bing Li, and Toshikazu Miyata for helping to make data collection possible. This work was supported by startup research funds from the Princeton Neuroscience Institute to J.G., and also based upon work supported by the National Science Foundation under CAREER Award 2337373 to J.G., and is the result of funding in part by the National Institutes of Health (NIH) through the National Eye Institute award number 5R01EY036881.

## Author contributions

Conceptualization, analysis, visualization, and manuscript preparation performed by E.D. & J.G.

## Data, Code, and Materials Availability

Code and data for reproducing figures can be found at: https://github.com/GitEdaniel/ASL-visual-categories-fmri

## Materials and Methods

### Participants

Data were collected from 33 participants: 11 Deaf signers (DS; learned to sign in childhood; 8 Females, mean age=46), 11 hearing sign language experts (HS; licensed interpreters/educators with >10y experience, learned to sign in college; 11 Females, mean age=47), and 11 controls with no sign language experience (C, 7 Females, mean age=45). All participants had normal or corrected-to-normal vision, and provided informed, written consent to participate in the experiment. Procedures were approved by the Princeton Internal Review Board on human subjects research (protocol 13074). Deaf signers were recruited from the Deaf community as individuals who were congenitally deaf and learned American Sign Language (ASL) in childhood. To recruit a group of hearing participants with expertise in sign language, we had two criteria. First, we sought to recruit participants who became fluent in ASL in adulthood. Second, while it is difficult to match the life-long experience with ASL present in Deaf signers, we wanted to ensure that expertise was standardized, and a long term experience. To that end, we recruited hearing sign language experts with over 10 years of signing experience, holding an external validation of fluency-either (1) Registered, Certified American Sign Language-English Interpreters (Registry of Interpreters for the Deaf; RID), or (2) holding an Educational Interpreter Performance Assessment (EIPA) Test score of 3.5 or higher, surpassing the most recent New Jersey Educational Interpreter Requirements.

### Data Acquisition

Magnetic resonance imaging (MRI) data were collected at the Scully Center for the Neuroscience of Mind and Behavior at the Princeton Neuroscience Institute. Participants completed data collection across two sessions, the first of which were used in the present analyses and described here. Data were collected on a Siemens 3-Tesla Skyra system, each participant completed structural scans, a visual category localizer, and a pRF Mapping Experiment. Following the MRI recording, participants completed an eyetracking experiment in an adjacent behavioral experiment facility.

Structural scans included a T1-weighted (voxel size 800μm3, TR=2.4s, TE=0.002s, Flip Angle = 8 degrees) and a T2-weighted (voxel size 800μm3, TR=3.2s, TE=0.565s, Flip Angle = 120 degrees) volume collected for each participant. These images were aligned and processed through FreeSurfer to reconstruct the cortical surface, with the T2-weighted image providing additional information for accurate reconstructions of the pial surface. Functional MRI scans comprised 72 slices acquired using a multiplexed echo planar imaging (EPI) sequence (multiband acceleration factor: 3, voxel size: 2.5mm isotropic), with repetition time (TR) = 2s, echo time (TE) = 31ms, and flip angle (FA) = 80°.

Eyetracking data was collected using an SR Research EyeLink 1000 Plus system. All participants used a chin rest with forehead mount to minimize head motion. The width of the monitor was 595mm and the height was 334mm, with a display resolution of 2560 pixels by 1440 pixels. The monitor was 895mm from the viewer’s eye. Participants’ eyes were calibrated using a 5-point calibration (four corners, one center) before stimulus presentation. Eye gaze position was recorded at a frequency of 1000Hz.

### Accessibility Measures

Communication with the participant during MRI scans is usually performed using an audio system. To ensure smooth communication with Deaf participants, scans were accompanied by a certified interpreter. Instructions were given before entering the MRI. During the scan, the interpreter remained in the control room and communicated with the participant through a bi-directional video system. In the control room, the in-scanner cameras allowed the interpreters to see the participants’ hands. The participant could see the interpreter on the screen in front of them through a video chat system to monitor for hand signs. The participants were asked to stay as still as possible, so the communication system was in place for reminders about task procedures, and to provide the ability of the participant to communicate questions or discomfort. The collection protocol for Deaf participants was developed with a Deaf collaborator to ensure our scans are comfortable and accessible, and approved by the Princeton Internal Review Board on human subjects research.

### Visual Category Localizer Task

Participants completed four runs of a visual category localizer (Stigliani et al., 2015). During each of four runs (209s per run) participants fixated on a central dot as visual stimuli were presented in 4-sec blocks. Each block consisted of eight images presented at a rate of 2Hz, from a single visual category out of seven in total: houses, hands, feet, faces (adult + child), full bodies (headless), pseudowords, and cars (Fig. 1a). Limb stimuli always included the digits. Each image was tightly cropped and overlaid on a textured grayscale background produced by phase scrambling a stimulus from another category, where an additional baseline condition consisted of a scrambled background with no overlaid image (Stigliani et al., 2015). Images subtended a visual angle of 7.125° centered on the fovea and were presented with PsychoPy (Peirce et al., 2019). To ensure attention to the stimuli, participants completed a 2-back task, where they were instructed to press a button whenever they detected that an image repeated itself with exactly one intervening image between repeats.

### Preprocessing of Visual Category Localizer fMRI Data

Preprecessing was performed in each individual’s native brain space, with no spatial smoothing. Data was preprocessed based on the HCP minimal preprocessing pipeline for motion-correction, slice-timing correction, and phase-distortion correction using Topup (Andersson et al., 2003; Smith et al., 2004). Resampling voxel time-series data to cortical vertices and general linear model (GLM) analyses were processed in each individual’s native brain space, through FsFast (FreeSurfer Functional Analysis Stream; https://surfer.nmr.mgh.harvard.edu/). An inflated cortical surface reconstruction was used for each participant and functional data were projected onto this cortical surface in which voxel time-series data were resampled to each cortical vertex. Using a GLM, we fit predictors to the data and estimated within each vertex the response amplitudes for each condition (betas), and residual variance of each vertex’s time course. The beta weights and residual variance from the GLM were used to generate contrast maps comparing responses of one visual category compared to another as described in the main text. The resulting contrast maps were then transformed into Freesurfer’s cortical average space for group analyses.

### Anatomical regions of interest (ROIs)

We focused our analyses to two main regions of interest: ventral surface and lateral surface in order to separate the ventral and lateral visual streams. Within the ventral surface, we further parcelated the cortex into the major anatomical landmarks: the collateral sulcus (CoS), the medial Fusiform gyrus (med-FG), the lateral Fusiform (lat-FUS), and the occipitotemporal sulcus (OTS), as illustrated in Figure 1. The lateral surface was parcelated into two components-the lateral occipitotemporal cortex (LOTC) and the superior temporal sulcus (STS).

### Identifying category selective regions

Category-selective regions of interest (ROIs) were defined using contrast maps derived from the visual category localizer experiment, in FreeSurfer’s average cortical space. Statistical contrasts of the category of interest > all other stimuli were thresholded at T-values greater than 3. For hand-selective regions, we used the statistical contrast of [hands > non-limb stimuli]. On the ventral surface, this included the hand-selective regions on the occipitotemporal sulcus (OTS-hand). On the lateral surface, we defined two ROIs: the lateral-occipital temporal cortex (LOTC), and the Superior Temporal Sulcus (STS), based on anatomical landmarks. LOTC-hand aggregated the hand-selective patches on the inferotemporal gyrus (ITG-hand), lateral occipital sulcus (LOS-hand), and middle temporal gyrus (MTG-hand) (Weiner & Grill-Spector, 2011). For face-selective regions, we used the contrast [faces > all other stimuli]. On the ventral surface, this included face-selective regions on the inferior occipital gyrus (IOG-faces), the posterior Fusiform (pFus-faces), the middle Fusiform gyrus (mFus-faces). On the lateral surface, this included selectivity for faces on the posterior branch of the superior temporal sulcus (pSTS-faces). For localization of the place-selective regions, we used the contrast [houses > all other stimuli]. On the ventral surface, this included the place-selective region within the collateral sulcus (CoS-place). On the lateral surface, this included the place selective region on the lateral occipital sulcus (LOS-place).

### Identifying regions of category overlap

To assess how face, hand, or house selectivity may overlap in lateral or ventral temporal cortex, we produced additional contrast maps similar to the procedure described above with a modification that would allow us to define each category-selective region while explicitly allowing for overlap of category-selective regions. To this end, we produced contrast maps for each category that excluded the other categories from the contrasting group. For example, we derived a face-selective contrast that excluded limbs in the contrasting stimulus group, as well as a hand-selective contrast map that excluded faces from the contrasting stimulus group, in order to identify cortex that is responsive to faces and hands (the conjunction of these two maps). Specifically, we contrasted each category with all inanimate categories [face > inanimate stimuli] and [hand > inanimate stimuli] to allow identification of shared face-hand (animate) representations. On the ventral surface, face and hand regions neighbor on the transition between the predominantly face selective fusiform gyrus (FG), and the predominantly limb-selective anterior Occipital-Temporal Sulcus (OTS). On the lateral surface, hand regions are interdigitated with face-selective regions (pSTS-faces and IOG-faces) surrounding motion-selective area hMT+ on the ascending limb of the posterior inferior temporal sulcus (Weiner & Grill-Spector, 2011). These hand-selective regions on the lateral surface also neighbor the LOS place-selective region located just posterior to the LOS-hands region.

### Statistical Analyses

To test for statistical significance of group differences across our dependent measures we used nonparametric bootstrap resampling with replacement across 1000 iterations, with two tailed 95% confidence intervals. In each iteration, we first calculated the bootstrapped statistic (S*) within each group. Then, for each iteration, we and then subtracted S* between groups to generate a distribution of group differences 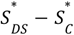 for comparing deaf signers to controls, and 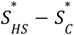 to compare hearing signers to controls), and quantified the effect size per iteration as 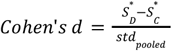 where 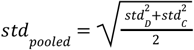. The within-iteration Cohen’s d values were then averaged over iterations to generate a bootstrapped estimate of effect size. Finally, we compared our group-difference bootstrapped-distributions to zero, quantifying the confidence intervals (CI) and p-values under the null hypothesis that there are no group differences 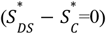. The P value was defined as the proportion of 1000 iterations that were of the opposite sign as the group difference, doubled for a two-tailed test, quantified as follows:

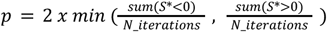

To test for main group effects and interaction effects, Linear Mixed Effects Models were used with subjects included as a random effect, implemented in Python using the statsmodels Module and smf.mixedlm function, complemented by a Wald Chi-Squared test to quantify the main fixed effects (Seabold & Perktold, 2010).

### Distance maps for estimating functional ROI shifts relative to anatomical landmarks

To quantify the distance of the functional ROIs from anatomical landmarks in ventral temporal cortex (VTC), we first labeled the anatomical structures of interest on the freesurfer average brain surface. We focus on the anterior border of the VTC for measuring anterior-posterior shifts, and the Collateral Sulcus (CoS) to measure medial-lateral shifts. We used the freesurfer function “mris_distance_transform” to create a distance map from each of the anatomical ROIs (Fig 4b), where the values denote the minimum distance in millimeters of each vertex to vertices within the anatomical reference ROI. Then, within each participant, visual category, and hemisphere, we quantified the average value in the intersection of the functional ROI and the distance map, quantifying the mean distance of the functional ROI from the anatomical landmark. These distances were then bootstrapped (N = 1000 iterations, with replacement, two tailed 95% confidence intervals) within each group, and compared between groups (Fig 4c, S4).

### Centroid analysis of functional ROI movement

To estimate how the centroid of a given category-selective region compares in spatial position between two groups, we employed a bootstrap (N = 500 iterations) approach to derive both the distance and angle of displacement between two functionally-defined ROIs. We compare the spatial location of face-selective region in the right hemisphere ventral temporal cortex located on the middle fusiform gyrus (mFus-faces) and the hand-selective region located in the occipitotemporal sulcus in the left hemisphere (OTS-hands). On each iteration, N = 11 participants are drawn with replacement from each group (Controls, Deaf Signers, Hearing Signers). To get a sense for how the distance of Signer groups compares to variability in Controls, an additional Control group is drawn with replacement on each iteration. For each bootstrapped group, unthresholded contrast maps from each participant are transformed to the FreeSurfer average cortical space using cortex-based alignment before averaging across participants within each group. Contrast maps are smoothed (FreeSurfer FWHM = 7) and then masked within a region surrounding the middle Fusiform gyrus or the middle OTS before thresholding maps at the 90th percentile to derive a group-average estimate of mFus-faces or OTS-hands ROIs. The centroid of each thresholded ROI is derived as the vertex that satisfies the condition of having the minimal total distance to all other vertices, and the distance between each group’s centroid is calculated using Dijkstra’s algorithm to derive the shortest path along edges connecting cortical vertices. The white matter surface was used to derive the distance, in millimeters, between a given pair of centroids. Distance in the Control group was counted as negative if the distance of the second Control group was further than the first Control group from Deaf signers, indicating it moved away. For Hearing Signers, if the distance between Hearing Signers and Deaf Signers was larger than the mean distance between Deaf Signers to Controls and Hearing Signers to Controls, this distance was counted as negative to indicate that the Hearing centroid had move away from the Control centroid in a direction distinct from Deaf signers. Otherwise it was counted as positive to indicate the Hearing centroid was displaced from the Control centroid in a direction similar to Deaf signers. On each iteration, we also derive an angle describing the vectors of displacement centered at the first Control group centroid and terminating at: Deaf Signers centroid, Hearing Signers centroid, or the second Control group centroid. The inflated cortical surface was used to derive vector angles given that ventral temporal cortex, on the highly inflated FreeSurfer average surface, is a nearly-flat plane where vectors are assumed to be largely parallel or “in-plane”. To obtain a standardized reference direction, we computed a vector pointing in the posterior-to-anterior direction, parallel with the MFS by establishing a vector connecting the posterior MFS to the anterior MFS. As an example, on a given iteration, we derive the vector between the first Control group and Deaf Signers (vector a) and compute the angle formed between this and the vector parallel with the MFS (vector b), given by *θ* = cos^−1^ 1(a ⋅ b / *∥* a *∥ ∥* b *∥*). The resulting angle was converted from radians to degrees for visualization. This operation is repeated for Hearing Signers relative to the first Control group, and also for the second Control group to estimate how the Control group displaces around itself as a function of bootstrapping.

### Intersection with independent atlas of category-selective regions in ventral temporal cortex

To compare the topographic location of the observed functional ROIs to an independent control dataset, we compared our selectivity maps to functional regions defined in the visfAtlas, which defines the probabilistic location of category-selective regions in human occipito-temporal cortex (Rosenke et al., 2021). While this atlas was built from localizer data that used very similar but slightly different stimulus categories (e.g., they define a body-selective regions in the OTS rather than a hand-selective region), it nonetheless serves as a useful independent reference to which all groups can be directly compared. For each participant, we quantified the number of vertices included within, and those outside of, the expected functional region of the OTS-body and the Fus-faces functional regions as defined by the independent probabilistic atlas. To test for group differences, we used a Linear Mixed Effect Model of the following form:

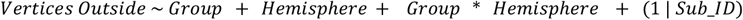

We then repeated the displacement vector analysis as detailed in the previous section, now using the centroid of the independently derived visfAtlas regions as reference points.

### Hemispheric Biases

To test for changes in hemispheric biases, we computed the Left:Right (L/R) ratio by dividing the value of interest in the left by the sum over both hemispheres, such that 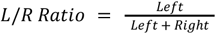. This ratio was quantified within each subject, visual category, and ventral/lateral surface. An L/R ratio<0.5 indicates Right hemisphere dominance, and L/R>0.5 indicates Left dominance (Fig. 5X). To test for group differences in the hemispheric biases, a Linear Mixed Effects Model was used, with the following form:

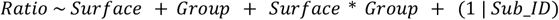

To test the statistical significance of the hemispheric biases of the different visual categories, a bootstrap analysis was performed similar to previous steps. The bootstrapped Ratios were then compared to 0.5 (null hypothesis: no hemispheric bias), with median ratios, confidence intervals, P values and bootstrapped effect sizes reported in the main text.

### Eye Tracking Experiment

Participants viewed 25 video clips, each 30-s in duration, selected from different Hollywood films (Costela & Woods, 2019). Movie clips were selected such that they included dynamic social stimuli, including people (with visible faces and bodies) either alone or engaging in social interactions, with other objects and a background scene present (Fig 5b) to allow participants the option to fixate on a range of animate and inanimate stimuli in any given clip. Participants were instructed to watch the clips as they normally would when enjoying a movie. Importantly, to keep the perceptual experience purely visual and consistent across groups, no sound was included during movie stimulus presentation, and eyetracking occurred outside the scanner. Participant head motion was controlled using a table-mounted chin rest.

### Eye Tracking data processing

To quantify viewing biases, eyetracking data was downsampled to 2Hz, and filtered to include only x,y eye positions during fixations (excluding saccades). Within each participant, we quantified the raw number of fixations made in each half of the stimulus (left, right, upper, lower), alongside the average x and y positions, and standard deviation on the horizontal (x) and vertical (y) axes. The spatial viewing biases were defined within each participant as the ratio of the number of fixations made to the one side of the visual stimulus normalized by the number of fixations to both sides, where:

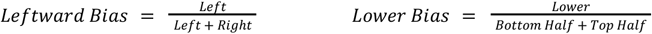

To quantify fixations made at different visual categories, movie stimuli were downsampled and individually labeled such that pixels containing faces, bodies, or background scenes could be differentiated. For this analysis, a subset of frames were extracted from each movie clip at a constant rate of 0.2 Hz, resulting in six frame samples per clip, for a total of 150 frames. The same frames were analyzed across all participants. Face and body stimuli were hand labeled using rectangles that subtended the vertical and horizontal extent of the pixels occupied by the face or body, with all other pixels defined as the background scene (Daniel-Hertz et al., 2025). Then, we computed the ratio of fixations on each visual category (face, body, scene) within-subject, by dividing the number of fixations on a single category by the overall number of fixations each participant made.

### Eye Gaze Fixation Biases

To compute the consistency of fixation biases across the different movie clips, we calculated the average x,y eye-gaze position across participants within each group, for each movie clip. We then measured the shift in the x (Δ*x* = *x* − *x*), and y positions (Δ*y* = *y*_2_ − *y*_1_), which allowed us to identify the vector that connects the group means for each clip, and quantify the euclidean shift 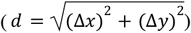 as well as the angle (Θ = *arctan*2(Δ*y*, Δ*x*)) between the group centers. The average angle across the n=25 movie clips was defined as: 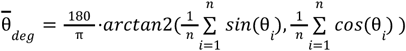. To measure the consistency of the vector directions, we used the Rayleigh Test of Uniformity, testing whether our vectors come from a uniform distribution with no preferred direction, or if there is consistent directionality between our experimental and control group means. Rayleigh’s R was defined as 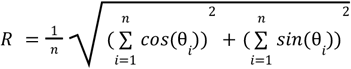 with R ranging between 0 and 1. The circular standard deviation (‘spread’) of the vectors was then calculated as: 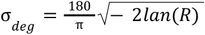 . Finally, to test the statistical significance of this directional shift, we used a spatial permutation test (paired, two-tailed, 5000 permutations), randomizing the direction of displacement (by multiplying the vector’s direction by either -1 or 1, randomly) to build a null distribution from the n=25 vectors (one vector per movie clip, connecting the group means). The P value was computed as the proportion out of the 5000 permutations in which the null distribution’s euclidean shift was equal to or larger than the observed shift.

To better visualize the spatial relation between the group gaze centers across the different movies, we centered the directional vectors such that the control group centers are at the origin (0,0) and the experimental group centers (Hearing or Deaf Signers) are positioned relative to this centering, maintaining the x,y axis units as pixels (Fig 5d, bottom row). We then quantified a 2D 95% confidence ellipse around the group average displacement, visualized such that we can see whether it overlaps with the origin (0,0) where the control group is centered. To this end, we bootstrapped the average x,y position of the spatial shift. On each iteration (1000 iterations total), N = 25 x,y positions of experimental group centers (Hearing or Deaf Signers, centered relative to Controls) from the different movie clips were drawn with replacement, and averaged to create a bootstrapped distribution of the spatial shift compared to controls. With this bootstrapped distribution of x,y positions, we could calculate the standard deviation of each variable (σ_*x*_, σ_*y*_), their covariance(σ _*xy*_), and Pearson Correlation. The sample covariance was defined as 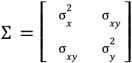, and the Pearson Coefficient as 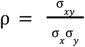. This allowed us to derive a standardized unit ellipse based on our data’s correlation dynamics, where 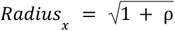 and 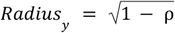. The ellipse was then centered around the data’s mean and scaled to correspond to the standard deviations (*σ*_*x*_, *σ*_*y*_), multiplied by a scaling factor to implement a ∼95% confidence interval for a 2D Gaussian distribution (2.447 std). This allowed us to visualize the 95% confidence interval of the spread of the displacement of the x,y positions, and see whether this ellipse includes the origin (0,0) where the control group is centered.

### LDA to test group separability based on viewing metrics

To test whether the experimental groups can be distinguished based on their viewing patterns, fixation metrics were entered into a linear discriminant analysis (LDA) which seeks to identify a linear combination of features to separate our three experimental groups. We used two LDs and plotted the distribution of participants against the two linear discriminant (LD) dimensions to visualize the group separability (**Fig 5E**). To quantify this, we computed the Silhouette Score (implemented with Python/sklearn.metrics/silhouette_score), which measures how similar each data point is to its own cluster (*a*_*i*_), compared to the nearest neighboring cluster (*b*_*i*_), providing a score between -1 and +1, where:

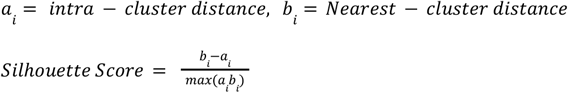

Such that positive scores indicate that the mean between-cluster exceeds intra-cluster distance.

#### Changes to variable coupling

The LDA analysis revealed the importance of the left-right and lower viewing biases for distinguishing group viewing behavior. Thus, we asked if the coupling between these two features is significantly different between groups by testing the linear relationship between them within each group, and comparing this relation across groups. To quantify the variable coupling within-group, we compute for each group the regression line using an Ordinary Least Squares (OLS) linear model (*Bias*_*Left*_ ∼*Bias*_*Lower*_), and report the slope (β), the P-value of the slope, and the variance explained by the model (*R*^2^). We then compare the difference in slopes (Δβ) across groups by running an interaction OLS model (*Bias*_*Left*_ ∼*Bias*_*Lower*_ *x Group*) to test whether the difference in slopes is statistically significant, reporting the P-value of the interaction. Finally, we compute the Bayes Factor (*BF*_10_) to quantify the strength of the evidence of the interaction effect, by comparing the Bayesian Information Criterion (BIC) of the interaction model with a null model that assumes no interaction, where groups share the same slope (*Bias*_*Left*_ ∼*Bias*_*Lower*_ + *Group*). The Bayes Factor is defined as 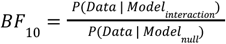, and was estimated as 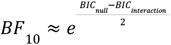.

Where possible, all individual data points were shown directly (e.g., swarm plots) or shown within visual summary illustrations (e.g., violin plots). Data analyses and visualizations were made using custom-built scripts in MATLAB (The Mathworks, Natick, MA), as well as using Python NumPy (Harris et al., 2020), Matplotlib (Hunter, 2007) and Seaborn (Waskom, 2021) packages.

Sample sizes are similar to those reported in previous work (Cardin et al., 2013; Gomez et al., 2019; Letourneau Mitchell, 2011; Sadato et al., 2004).

**Figure S1:**
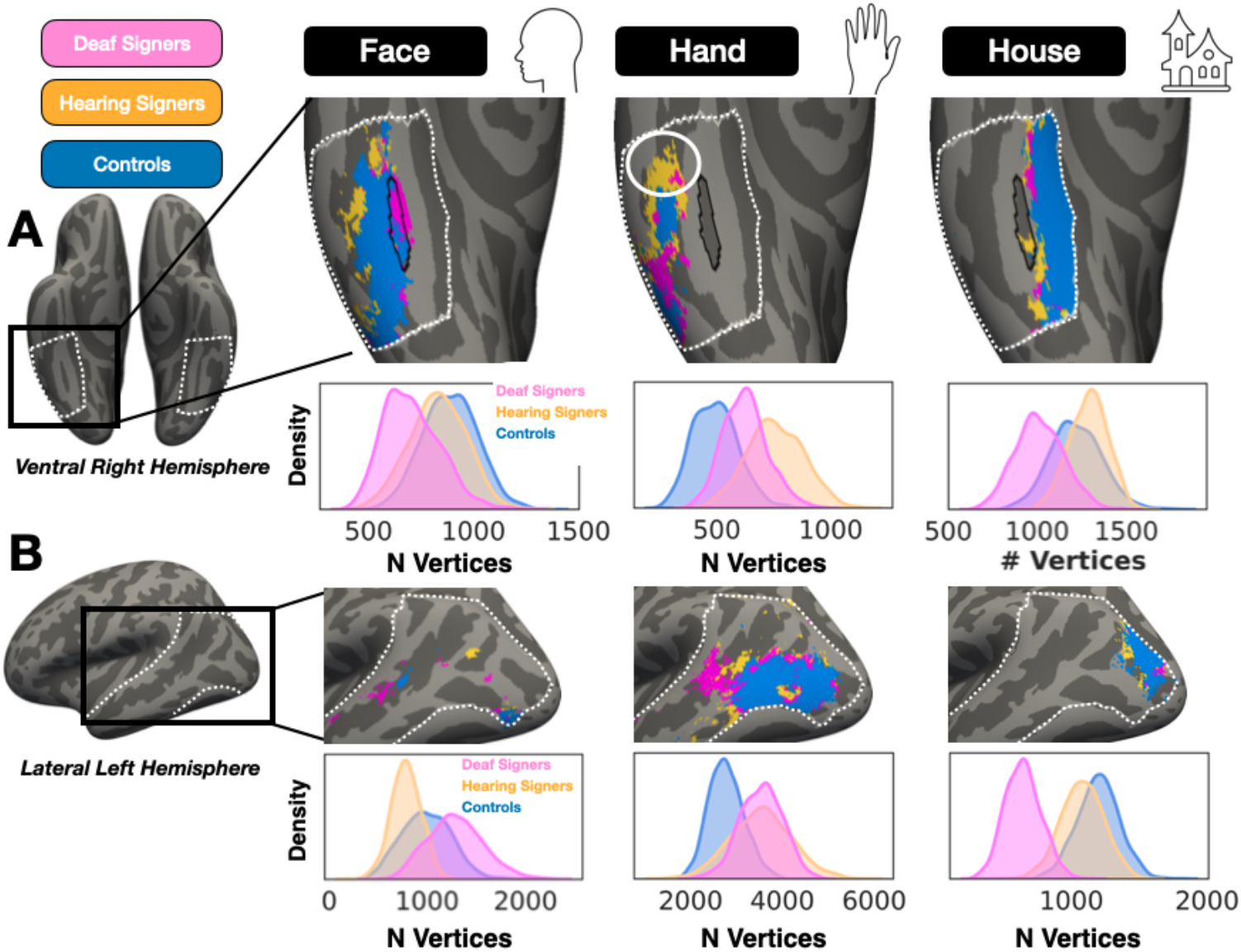
Expansion of hand representations in Deaf and hearing signers, in ventral right and lateral left hemispheres. **(A)** Face, hand, and house selective regions in ventral temporal cortex (dashed white border). Contrast maps were thresholded at T>3 and binarized within individuals, then summed across individuals per group and thresholded at minimum of 4 participants to create the group-level category selective region. The mid-Fusiform sulcus (MFS) is outlined in black as an anatomical reference. Histograms depict bootstrapped (n = 1000) distributions of surface area (number of vertices in average cortical space) in each group, drawn with replacement. **(B)** Category-selective regions of similar categories but within left lateral visual cortex (dashed white border).

**Figure S2:**
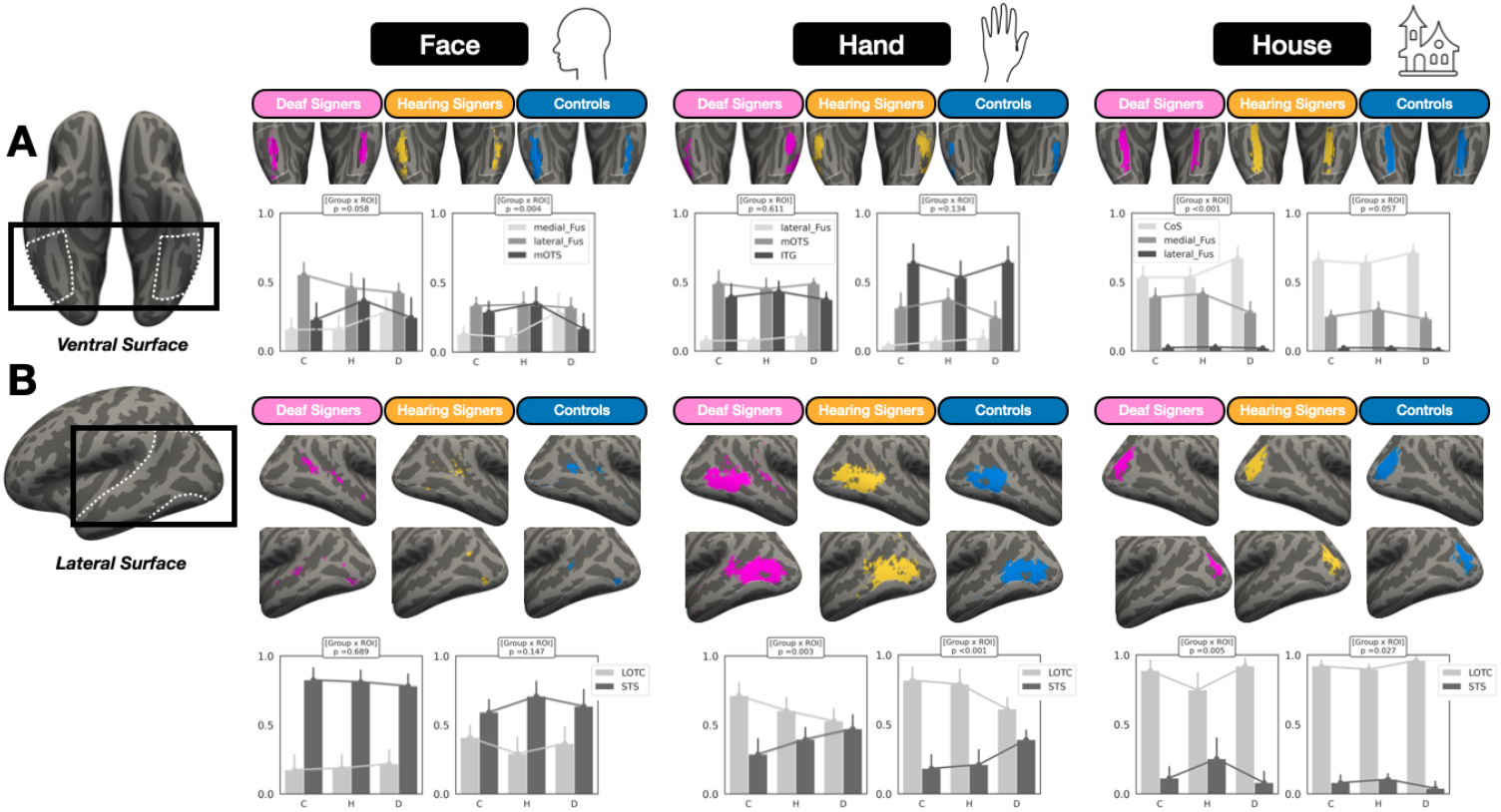
Topographic analysis of single-subject category-selective regions by anatomical region: **(A)** In ventral temporal cortex (white dotted line), the fraction of total vertices selective for a given category (thresholded at T > 3) in a given participant binned by anatomical structure. C = Controls, H = Hearing signers, D = Deaf signers. Anatomical structures of interest included: medial Fusiform, lateral Fusiform, middle occipitotemporal sulcus (mOTS), inferior temporal gyrus (ITG), and collateral sulcus (CoS). Bar height denotes mean value across participants within a group, error bar denotes bootstrapped 95% confidence intervals around the group means (n = 1000). For each category, the left skyline plots are data from the left hemisphere, and the plots on the right are from the right hemisphere. **(B)** Similar to the previous panel but for vertices in lateral visual cortex (white dotted line) within anatomical structures of lateral occipitotemporal cortex (LOTC) and the superior temporal sulcus (STS). Related to Figure 4A.

**Figure S3.**
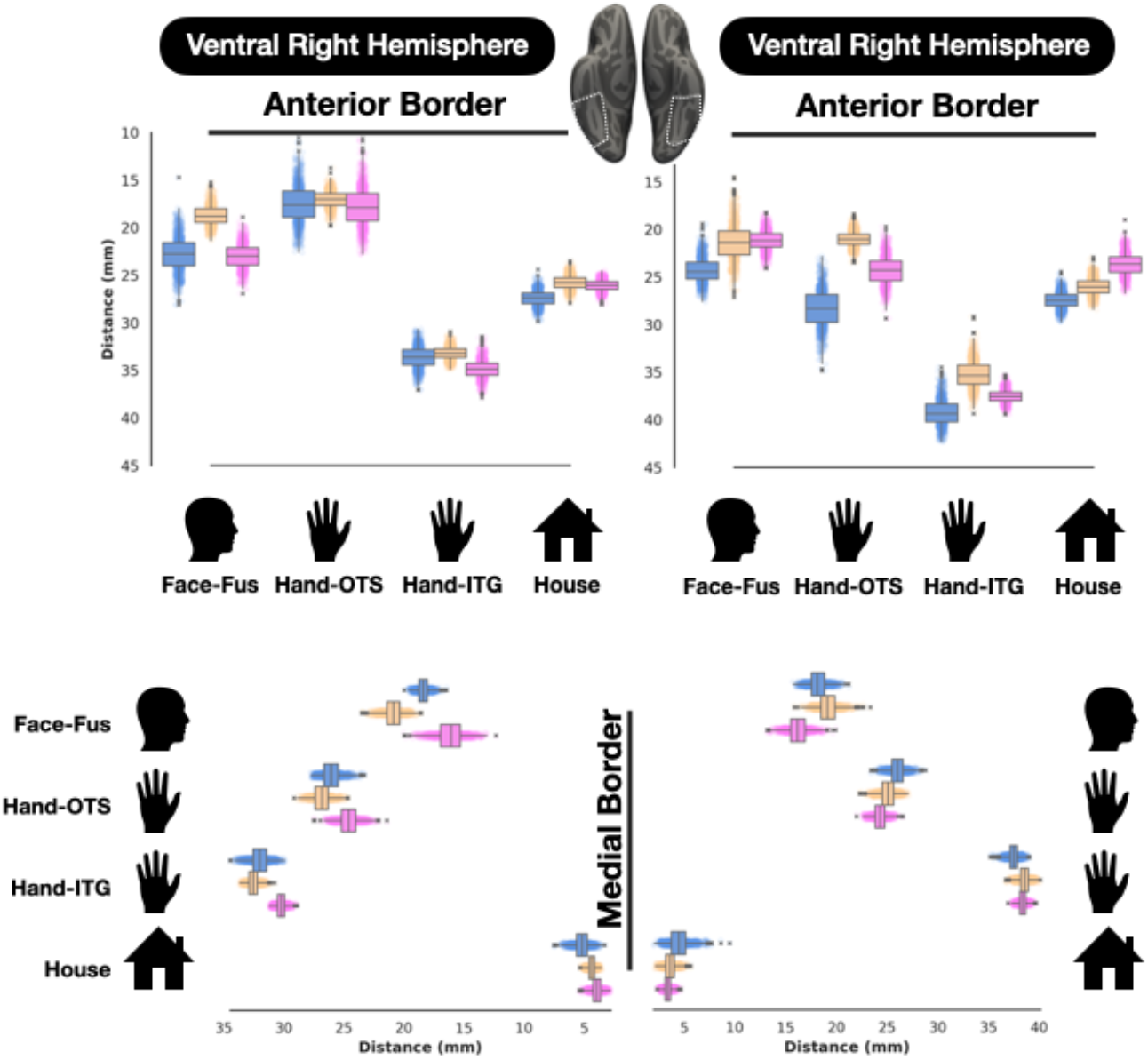
Sign language experience shifts visual map boundaries. Box plots depicting bootstrapped mean distance values of category selective regions from anatomical landmarks, similar to Figure 4C. Plotted here are: face-selective vertices on the fusiform gyrus (Fus), hand-selective vertices in the occipitotemporal sulcus (OTS), hand-selective vertices on the inferior temporal gyrus (ITG), and house-selective vertices in the collateral sulcus (CoS). Distances computed from either the anterior (top row) or medial (bottom row) anatomical structure of interest, as shown in main Figure 4B. Because hand-selectivity was expanded and more medially located in signers, the ITG-hand region had many vertices occupying the posterior aspect of the OTS. Rather than group these distinct vertices with the OTS-hands region, we quantify their position separately from OTS-hands given that they are functionally distinct regions.

## References

1. E. E. Meyer, M. Martynek, S. Kastner, M. S. Livingstone, M. J. Arcaro, Expansion of a conserved architecture drives the evolution of the primate visual cortex. Proc Natl Acad Sci U S A 122, e2421585122 (2025).

2. M. J. Arcaro, M. S. Livingstone, A hierarchical, retinotopic proto-organization of the primate visual system at birth. Elife 6 (2017).

3. D. Maurer, Critical periods re-examined: Evidence from children treated for dense cataracts. Cogn. Dev. 42, 27–36 (2017).

4. K. Srihasam, J. L. Vincent, M. S. Livingstone, Novel domain formation reveals proto-architecture in inferotemporal cortex. Nat. Neurosci. 17, 1776–1783 (2014).

5. K. Grill-Spector, K. S. Weiner, The functional architecture of the ventral temporal cortex and its role in categorization. Nature Reviews Neuroscience 15, 536–548 (2014).

6. J. Gomez, V. Natu, B. Jeska, M. Barnett, K. Grill-Spector, Development differentially sculpts receptive fields across early and high-level human visual cortex. Nat. Commun. 9, 788 (2018).

7. S. Dehaene, L. Cohen, J. Morais, R. Kolinsky, Illiterate to literate: behavioural and cerebral changes induced by reading acquisition. Nat Rev Neurosci 16, 234–244 (2015).

8. S. Dehaene, F. Pegado, L. W. Braga, P. Ventura, G. Nunes Filho, A. Jobert, G. Dehaene-Lambertz, R. Kolinsky, J. Morais, L. Cohen, How learning to read changes the cortical networks for vision and language. Science 330, 1359–1364 (2010).

9. M. J. Arcaro, P. F. Schade, J. L. Vincent, C. R. Ponce, M. S. Livingstone, Seeing faces is necessary for face-domain formation. Nat. Neurosci. 20, 1404–1412 (2017).

10. M. Nordt, J. Gomez, V. Natu, A. A. Rezai, D. Finzi, H. Kular, K. Grill-Spector, Cortical recycling in high-level visual cortex during childhood development. Nat Hum Behav 5, 1686–1697 (2021).

11. K. Emmorey, R. Thompson, R. Colvin, Eye gaze during comprehension of American Sign Language by native and beginning signers. J Deaf Stud Deaf Educ 14, 237–243 (2009).

12. S. M. Letourneau, T. V. Mitchell, Gaze patterns during identity and emotion judgments in hearing adults and deaf users of American Sign Language. Perception 40, 563–575 (2011).

13. D. Bavelier, M. W. G. Dye, P. C. Hauser, Do deaf individuals see better? Trends Cogn Sci 10, 512–518 (2006).

14. L. Kliger, G. Yovel, Distinct Yet Proximal Face- and Body-Selective Brain Regions Enable Clutter-Tolerant Representations of the Face, Body, and Whole Person. J Neurosci 44 (2024).

15. K. S. Weiner, K. Grill-Spector, Not one extrastriate body area: using anatomical landmarks, hMT+, and visual field maps to parcellate limb-selective activations in human lateral occipitotemporal cortex. Neuroimage 56, 2183–2199 (2011).

16. K. S. Weiner, K. Grill-Spector, Neural representations of faces and limbs neighbor in human high-level visual cortex: evidence for a new organization principle. Psychol. Res. 77, 74–97 (2013).

17. K. Grill-Spector, The neural basis of object perception. Curr Opin Neurobiol 13, 159–166 (2003).

18. M. Rosenke, R. van Hoof, J. van den Hurk, K. Grill-Spector, R. Goebel, A Probabilistic Functional Atlas of Human Occipito-Temporal Visual Cortex. Cerebral cortex (New York, N.Y. : 1991) 31 (2021).

19. B. Rossion, A. Lochy, Is human face recognition lateralized to the right hemisphere due to neural competition with left-lateralized visual word recognition? A critical review. Brain Structure and Function 227, 599–629 (2021).

20. M. Behrmann, D. C. Plaut, Hemispheric Organization for Visual Object Recognition: A Theoretical Account and Empirical Evidence. Perception 49, 373–404 (2020).

21. A. T. Smith, M. W. Greenlee, K. D. Singh, F. M. Kraemer, J. Hennig, The processing of first- and second-order motion in human visual cortex assessed by functional magnetic resonance imaging (fMRI). J. Neurosci. 18, 3816–3830 (1998).

22. E. Grossman, M. Donnelly, R. Price, D. Pickens, V. Morgan, G. Neighbor, R. Blake, Brain areas involved in perception of biological motion. J. Cogn. Neurosci. 12, 711–720 (2000).

23. L. Isik, K. Koldewyn, D. Beeler, N. Kanwisher, Perceiving social interactions in the posterior superior temporal sulcus. Proc Natl Acad Sci U S A 114, E9145–E9152 (2017).

24. D. Pitcher, L. G. Ungerleider, Evidence for a Third Visual Pathway Specialized for Social Perception. Trends Cogn Sci 25, 100–110 (2021).

25. F. M. Costela, R. L. Woods, A free database of eye movements watching “Hollywood” videoclips. Data in Brief 25, 103991 (2019).

26. E. Daniel-Hertz, J. K. Yao, S. Gregorek, P. M. Hoyos, J. Gomez, An Eccentricity Gradient Reversal across High-Level Visual Cortex. J Neurosci 45 (2025).

27. J. Gomez, M. Barnett, K. Grill-Spector, Extensive childhood experience with Pokémon suggests eccentricity drives organization of visual cortex. Nat Hum Behav 3, 611–624 (2019).

28. N. Hedger, T. Naselaris, K. Kay, T. Knapen, Vicarious body maps bridge vision and touch in the human brain. Nature 650, 173–181 (2026).

29. M. Coppola, K. Walker, Early language access and STEAM education: Keys to optimal outcomes for deaf and hard of hearing students. Educ. Sci. (Basel) 15, 915 (2025).

## References

Andersson, J. L. R., Skare, S., & Ashburner, J. (2003). How to correct susceptibility distortions in spin-echo echo-planar images: application to diffusion tensor imaging. NeuroImage, 20(2), 870–888.

Cardin, V., Orfanidou, E., Rönnberg, J., Capek, C. M., Rudner, M., & Woll, B. (2013). Dissociating cognitive and sensory neural plasticity in human superior temporal cortex. Nature Communications, 4(1), 1473.

Costela, F. M., & Woods, R. L. (2019). A free database of eye movements watching “Hollywood” videoclips. Data in Brief, 25, 103991.

Daniel-Hertz, E., Yao, J. K., Gregorek, S., Hoyos, P. M., & Gomez, J. (2025). An Eccentricity Gradient Reversal across High-Level Visual Cortex. The Journal of Neuroscience : The Official Journal of the Society for Neuroscience, 45(2). 10.1523/JNEUROSCI.0809-24.2024

Gomez, J., Drain, A., Jeska, B., Natu, V. S., Barnett, M., & Grill-Spector, K. (2019). Development of population receptive fields in the lateral visual stream improves spatial coding amid stable structural-functional coupling. NeuroImage, 188, 59–69.

Harris, C. R., Millman, K. J., van der Walt, S. J., Gommers, R., Virtanen, P., Cournapeau, D., Wieser, E., Taylor, J., Berg, S., Smith, N. J., Kern, R., Picus, M., Hoyer, S., van Kerkwijk, M. H., Brett, M., Haldane, A., Del Río, J. F., Wiebe, M., Peterson, P., … Oliphant, T. E. (2020). Array programming with NumPy. Nature, 585(7825), 357–362.

Hunter. (2007). Matplotlib: A 2D Graphics Environment. 9, 90–95.

Letourneau, S. M., & Mitchell, T. V. (2011). Gaze patterns during identity and emotion judgments in hearing adults and deaf users of American Sign Language. Perception, 40(5), 563–575.

Peirce, J., Gray, J. R., Simpson, S., MacAskill, M., Höchenberger, R., Sogo, H., Kastman, E., & Lindeløv, J. K. (2019). PsychoPy2: Experiments in behavior made easy. Behavior Research Methods, 51(1), 195–203.

Sadato, N., Yamada, H., Okada, T., Yoshida, M., Hasegawa, T., Matsuki, K.-I., Yonekura, Y., & Itoh, H. (2004). Age-dependent plasticity in the superior temporal sulcus in deaf humans: a functional MRI study. BMC Neuroscience, 5, 56.

Seabold, S., & Perktold, J. (2010). Statsmodels: Econometric and statistical modeling with python. Proceedings of the Python in Science Conference, 92–96.

Smith, S. M., Jenkinson, M., Woolrich, M. W., Beckmann, C. F., Behrens, T. E. J., Johansen-Berg, H., Bannister, P. R., De Luca, M., Drobnjak, I., Flitney, D. E., Niazy, R. K., Saunders, J., Vickers, J., Zhang, Y., De Stefano, N., Brady, J. M., & Matthews, P. M. (2004). Advances in functional and structural MR image analysis and implementation as FSL. NeuroImage, 23 Suppl 1, S208–S219.

Stigliani, A., Weiner, K. S., & Grill-Spector, K. (2015). Temporal Processing Capacity in High-Level Visual Cortex Is Domain Specific. The Journal of Neuroscience : The Official Journal of the Society for Neuroscience, 35(36), 12412–12424.

Waskom, M. (2021). seaborn: statistical data visualization. Journal of Open Source Software, 6(60), 3021.

Weiner, K. S., & Grill-Spector, K. (2011). Not one extrastriate body area: Using anatomical landmarks, hMT+, and visual field maps to parcellate limb-selective activations in human lateral occipitotemporal cortex. NeuroImage, 56, 2183–2199.

